# Modelling a rapid radiation of crown-group placentals

**DOI:** 10.64898/2026.08.21.746252

**Authors:** Mary Kate Branigan, Richard P. Mann, Graham E. Budd

## Abstract

The timing of the origins of the crown-group placental mammals has provided one of the classic battlefields in the long-running debate over when clades arise. Undoubted fossil crown-group placentals appear only in the Paleogene, but even so most molecular analyses, and many palaeontologists, have suggested their true origin is somewhere between 70-100 Ma. However, apart from the fact of the fossil record itself, there are several reasons to believe that the true origin is indeed post-Cretaceous, including consideration of the dynamics of stem and crown groups, which strongly favour crown-group origins to lie just after, and not just before, mass extinctions. Here we consider this “hard explosive” model in the light of the newly-developed “Covariant Evolutionary Tempo (CET)” model which allows diversification and molecular evolution rates to covary. It predicts “early bursts” in both lineage creation and molecular evolution at the base of major radiations which lead to highly unequally-sized clades; and an inheritance of rapid rates from this initial event by extant rapidly-evolving clades. We show that when the placentals are constrained to emerge after the K-Pg boundary, they indeed show elevated rates of both diversification and molecular evolution, which rapidly decline. Nevertheless, although elevated, these rates are comparable to the fastest rates seen in extant clades such as the rodents. In addition, the contiguous lineages leading from the origin to the rodents and other fast evolving clades also show elevated rates. These patterns suggest that not only is a Paleogene origin for the placental crown-group plausible, as fossil evidence suggests, but they also provide support for the CET model, which should be considered in other cases of pronounced fossil record/molecular clock mismatch.

## Introduction

Why are there no Cretaceous crown-group placental mammal fossils? Paleontological discoveries have in general provided a substantial framework for our understanding of the origin of clades, with the fossil record showing distinctive patterns in taxon shifts particularly after mass extinction events (Jablonski, 2001; McElwain & Punyasena, 2007). While the fossil record is typically considered a good representation of diversification and evolutionary patterns through time by palaeontologists, it only preserves a relatively small fraction of biodiversity due to factors such as preservation potential, diagenetic influence, and population size (Benton, 2003; Benton et al., 2000; Budd & Mann, 2018; Cooper et al., 2006; Foote et al., 1999; Foote & Sepkoski, 1999; Kidwell & Holland, 2002; Woolley et al., 2025). The incompleteness of the fossil record is often cited as a reason why it should not be used as an accurate depiction of the origin of clades, and rather, that tools such as molecular clocks estimate the age of clade origins more accurately and effectively (e.g. Carlisle et al., 2023). However, molecular clocks often considerably overestimate clade origins relative to the fossil record, which has led to a great deal of debate about why this might be the case.

Placental mammals are in particular one of the groups of major contention between palaeontologists and molecular clock proponents in terms of the origin of the crown group. The Cretaceous-Palaeogene (K-Pg) extinction at c. 66 Ma is marked by a fundamental shift in the fossil record where characteristic Mesozoic taxa such as dinosaurs abruptly disappear and are replaced by modern taxa such as crown-group placental mammals and birds. In particular, the rapidity of appearance of diverse placental groups in the Palaeogene has led to protracted discussion about what should be inferred about the true timing of the crown-group placentals origin. Although a straightforward reading of the fossil record might suggest an explosive origin of placentals directly after the boundary (Wible et al. 2007; O’Leary et al. 2013a), this interpretation has been vigorously contested (Springer et al. 2013), both on the grounds of the implied rapidity of the event, but also by various molecular clock studies, which have in recent years been joined by various attempts at modelling at how the fossil record itself is generated. In addition, reconstruction of the nature of the last common ancestral forms of crown group has suggested that it might be difficult to distinguish them from the stem group members, making correct phylogenetic assignment of fossils either side of the boundary problematic (e.g. Álvarez-Carretero et al., 2022; Archibald & Deutschman, 2001; Bininda-Emonds et al., 2007; Carlisle et al., 2023; Springer et al., 2019; Halliday et al. 2019).

The dispute over the timing of the radiation of the placental mammals has led to their possible time of origin being characterised by various different monikers, depending on whether the origin is considered to be well before, shortly before, or after the K-Pg boundary i.e. the long fuse, short fuse, and soft and hard explosive models respectively (e.g. Phillips 2016, Carlisle et al. 2023). The first two scenarios imply that stem-and crown-group placentals coexisted for a relatively long period of time within the Cretaceous, as well as that the pre-Paleogene crown group survived the mass extinction. Primary support for the short and long fuse scenarios has come from relaxed molecular clock studies which indeed typically infer the origin of placental mammals to lie within the Cretaceous (Foley et al. 2023; Álvarez-Carretero et al., 2022; Carlisle et al., 2023; dos Reis et al., 2014). However, despite this result, it remains problematic from the perspective of the fossil record, as no unequivocal evidence of crown-group placentals exists until the Paleogene (Halliday et al., 2017; de Vries & Beck, 2023). In addition, the characterization of these scenarios is at least partly suspect as they rely on a differentiation between “inter-” and “intra”-ordinal diversification, even though it is widely recognized that Linnean orders do not carry any particular importance apart from typically being large clades. Such a differentiation is sometimes reflected in calibration practice, so that in Phillips (2016) the superordinal groups are constrained to lie after the K-Pg boundary, and in Álvarez-Carretero et al. (2021), many of the orders are.

Despite this rather striking pattern, several attempts have been made to defend a Cretaceous crown-group placental origin from modelling the fossil record itself. These include a schematic parameterisation of the known record of primates to infer an origin in the late Cretaceous (Tavaré et al., 2002; Wilkinson et al., 2011; see Dos Reis et al., 2014), and the more recent “Bayesian Brownian Bridge” (BBB; Carlisle et al., 2023) of the entirety of placentals. Whilst we acknowledge the importance of modelling the fossil record, we note that all of these attempts are model-dependent (c.f. comments in Phillips, 2016), and stress again that there are no clear signs of Cretaceous crown group placental fossils. This point is reinforced by calculation of stem-and crown-group dynamics (Budd & Mann, 2020) which demonstrates that crown groups are unlikely to emerge just before a mass extinction, and very likely to emerge just after. In addition, if the placental crown group were to have emerged before the extinction, its diversity is likely to have been comparable to that of the stem group close to the end of the Cretaceous. This suggests that if crown-group placentals did emerge before the boundary, their fossil record should be comparable to that of the stem-group forms. Because the preservability of late stem and early crown-group mammals should also be comparable (Phillips, 2016), there seems no good reason for the numerous Cretaceous mammal sites not to preserve crown-group forms if present.

One further reason for thinking that the crown-group placentals could have emerged rapidly is provided by the newly-developed “Covariant Evolutionary Tempo” model (Budd & Mann, 2025), which allows rates of both diversification and molecular evolution to covary. In such a model, large clades are expected to have commenced with high evolutionary tempo, generating both a very large number of species and a high level of molecular change. This intense early period of change is compatible with a rapid crown-group placental radiation directly after the K-Pg boundary, but to conventional models of evolution, this amount of change would seem to imply an extended period of time. Another key prediction of the CET is that rapidly evolving species today in such clades are likely to have inherited their rates from the initial radiation, leaving a contiguous lineage of elevated rates of molecular evolution as a historical signature of the process.

Although the origin of the crown-group placentals in the Cretaceous has thus been described as being “settled” (Dos Reis et al., 2014), we would argue that the data at hand, plus our conceptual and theoretical considerations discussed above, actually point to the explosive model of crown-group placentals emerging after the K-Pg boundary being at least plausible. As a result, in this paper we show what implications such a model has for clock dating and the nature of the earliest crown-group placentals.

A previous attempt to show the effects of tight, late priors on the origin of the placentals (Phillips, 2016) placed Paleogene soft maximums on the ages of primates, rodents and bats for his main analysis, whilst placing a hard maximum at the K-Pg boundary origin of Afrotheria, Xenarthra, Laurasiatheria, and Euarchontoglires for his rates analysis, thus testing the “soft explosive” model. Here we examine trees and rates in the “hard explosive” model of constraining all crown-group placental mammals to emerge after the K-Pg boundary.

## Materials and Methods

In all of our analyses, we use PAML including BASEML and MCMCTree (Yang, 2007). Unless stated otherwise, we follow the procedure and settings of Álvarez-Carretero et al. (2021) in their initial analysis of 72 taxa.

In a previous attempt to infer placental origins using tight priors on placental origins (Budd & Mann, 2023) the branch lengths, gradient and Hessian used for the approximate likelihood calculation were estimated in BASEML using “method = 0”. On a dataset of this size, this algorithm may terminate before reaching the maximum likelihood estimates, so the Hessian was evaluated away from the optimum. We are grateful to Sandra Álvarez-Carretero and Mario dos Reis for drawing this to our attention. We have thus repeated the analyses using “method = 1”, which converges reliably for large datasets, and report the revised results below. The original placental mammal dataset of Álvarez-Carretero et al. (2021), including genomic data from 72 extant mammalian taxa, was used to conduct molecular clock analyses for the origin of the crown placental clade. Because calculations of the posterior can be very time consuming with large datasets like that used here, we have used the approximation method described by dos Reis and Yang (2011) which estimates the branch lengths then the gradient and Hessian for the branch lengths. Following Álvarez-Carretero et al. (2021), Baseml calculations of the Hessian for the dataset from the publication were used in MCMC calculations of both the prior and posterior. Alignment datasets as well as Hessian calculations and other relevant data from the analysis of Álvarez-Carretero et al. (2021) can be found at: https://doi.org/10.6084/m9.figshare.14885691. Each run was performed 4 times and examined for convergence.

We first re-ran the analysis of Budd and Mann (2023). As in this paper, all the calibrations and settings were as in Álvarez-Carretero et al., 2021, with the exception of the calibration on the crown-group placental node, which was set with a soft maximum age of 66 Ma to coincide with the K-Pg boundary. With this set of calibrations, the posterior for the age of the placentals was approximately 70 Ma with a 95% HPD of c. 74.2 Ma – 66.1 Ma, differing from our previous result, and is inconsistent with a post-Cretaceous placental origin. This result would seem to be in conflict with the maximum age set at 66 Ma in the calibration, but this difference is the result of a subtlety in how MCMCTree calibrations are typically set. In MCMCTree, fossils are used to introduce calibration ranges on nodes of a fixed tree. The calibration is a user-generated probability distribution of the true age of the node being calibrated. This can take a variety of forms, but the simplest and most widespread version is a uniform distribution from the age of the oldest fossil back to an age where it is considered that descendants of the node in question are unlikely to have evolved yet. In order to account for error in this assignment (for example, misidentification of the oldest fossil or choosing an erroneous maximum age) this uniform distribution is typically extended in both the younger and older directions, so that 2.5% of the total probability lies beyond each user-defined end of the calibration, in the form of an exponential distribution. Thus, although most of the calibration density lies within the original bounds, the calibration in theory allows ages older, or much younger, than the original uniform bound. When fossil calibrations are converted into effective priors in MCMCTree, there tends to be a bias towards older ages for deep nodes (Brown & Smith, 2018; Budd & Mann, 2023); the soft (rather than hard) maximum boundary allows both the effective prior for the placentals to be heavily peaked on, and indeed just before, the boundary. An important principle of Bayesian inference is that the effective priors (as opposed to the raw calibration densities) should represent our beliefs about the dating of nodes resulting from fossil analysis and before integrating molecular evidence; thus, our aim was first to establish priors that lie substantially within the Paleogene as a necessary condition for exploring the consequences of enforcing a post-boundary placental origin. Preliminary calibration tests were conducted with soft maximum calibrations for the base of the Placentalia crown group placed between 60 and 66Mya at 100ka intervals to test whether significant probability density of both the effective prior and posterior lay before or after the K-Pg boundary. These preliminary tests showed that as the soft maximum calibration age of the placentals decreased, both the effective prior (as expected) and the posterior also decreased linearly but at different rates, so that the effective prior and the posterior slowly converge. When the calibration maximum age reaches 64.5 Ma, the posterior already has its 95% HDP reaching into the Paleogene, and calibration of the placentals with a soft maximum age of 63Mya resulted in an effective prior and posterior estimate that both lay substantially after the K-Pg boundary (Fig 1). We thus selected 63Mya as a soft maximum age calibration which would place the origin of Placentalia in the Paleogene, in order to test the effect on rates along the tree when the molecular clock analysis is forced to consider a post K-Pg origin for the clade. The other placental calibration used, within the Cretaceous, was set to 80Mya to reflect the (posterior) results of Álvarez-Carretero et al. (2021) to provide a representative older prior dataset to compare against the younger results. Following Álvarez-Carretero et al. (2021), a relaxed correlated clock model was used with birth death process values for priors on nodes without fossil calibrations; µ and *λ* were set to 1 with a sampling fraction ρ of 0.1. We note that these are essentially arbitrary values but reflect those used in Álvarez-Carretero et al. (2021). Analyses were run using MCMCtree (Rannala, 2007; Yang & Rannala, 2006) in PAML 4.9j (Yang, 2007). Resulting rates of molecular substitution across the tree were then plotted along the tree as well as against each other for each of four partitions from the original publication along with an average of all partitions together with 95% HPD in R (v 4.5.3; R Core Team 2021).

**Figure 1.**
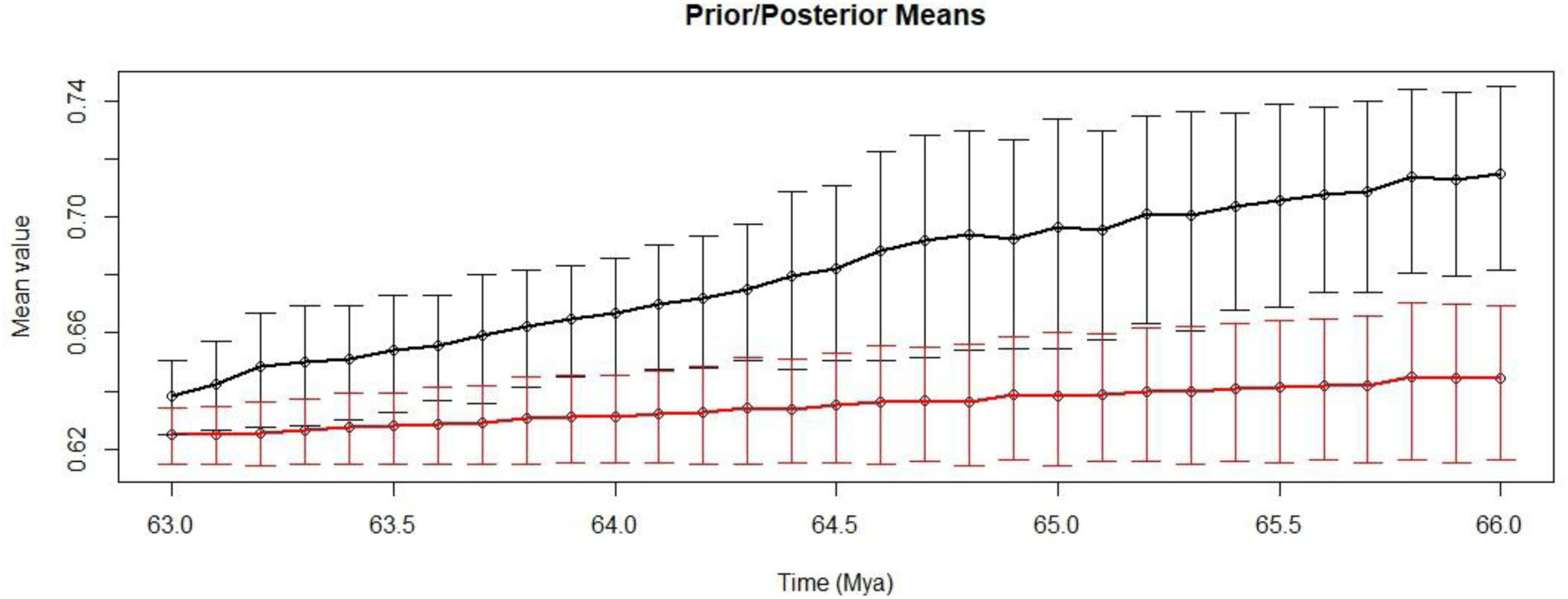
Plot of effective prior (red) and posterior (black) means and 95% HPD at 100ka intervals for the base of Placentalia. Note that as the soft maximum calibration age of the Placentalia node decreases, the two values converge and, as expected given the decreasing space between the soft maximum age and the calibration of younger nodes, the 95% HDP intervals also shrink (numbers given in Supplementary Table 1).

## Results

As the soft maximum age calibration for the base of Placentalia was decreased from 66 Ma to 63 Ma, the prior and posterior density distributions and their 95% HPD shifted towards each other, with younger calibration ages showing a definitive overlapping of the 95% HPDs while older calibrations fail to overlap at all (Fig. 1), thus demonstrating that the posterior remains sensitive to the prior across the range of interest.

Because one of the critiques of a Paleogene origin of placental mammals has been that it would imply “viral” rates of molecular change (Springer et al. 2013, Álvarez-Carretero et al. (2026)), we examined the posteriors of the rates implied by the study of Álvarez-Carretero et al. (2021). This study partitioned the molecular data into four partitions based on rates of change. We thus chose the fastest of these partitions to illustrate evolutionary rates in the most “extreme” case. For this partition, lineages through time see a sharp increase at the start of the Paleogene just after the K-Pg mass extinction event when the placental age prior mean is set to be after the K-Pg boundary (in this case, with the calibration soft maximum set at 63 Ma), in line with the hypothesis for crown emergence after the mass extinction as the fossil record suggests (Fig. 2). Lineages have a similarly sharp increase with the pre-K-Pg boundary calibration (80 Ma); however, this increase occurs prior to the mass extinction event (Fig. 3).

**Figure 2.**
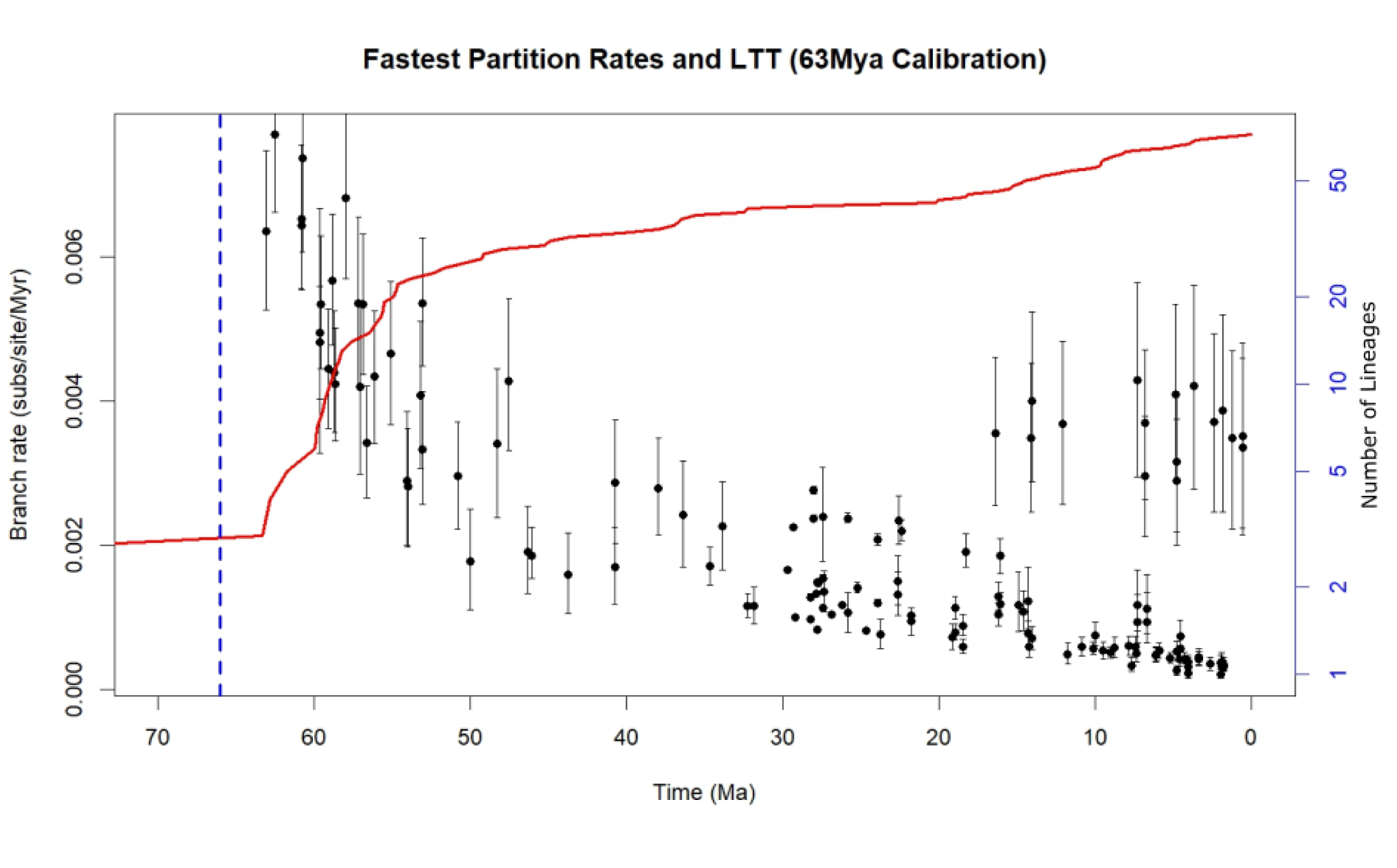
Rates of molecular change (black) along the branches of the Placentalia tree for the fastest partition with a 63Mya base calibration. Lineages through time are plotted in red, and the blue dashed line indicated the K-Pg boundary.

**Figure 3.**
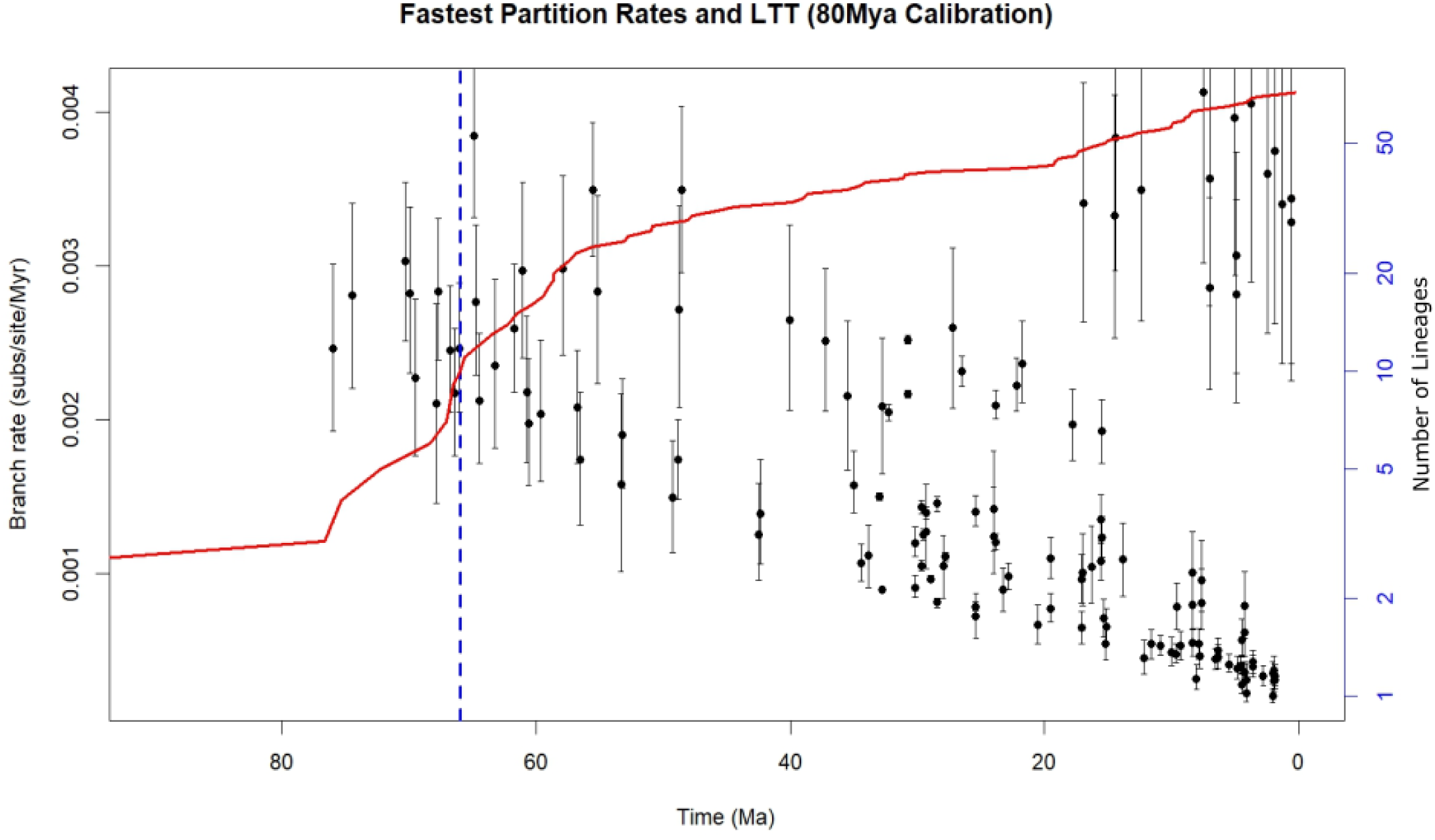
Rates of molecular change (black) with 95% confidence intervals along the branches of the Placentalia tree for the fastest partition with 80Mya base calibration. Lineages through time are plotted in red, and the blue dashed line indicates the K-Pg boundary. The crown placental node is located at c. 76.6Mya.

The substitution rates with the post K-Pg boundary calibration begin relatively high in both scenarios and rapidly decrease over time into the present, with some branch rates remaining higher closer to the present. The highest rates are approximately 0.006-0.007 substitutions per base pair per million years for the post KPg calibration and roughly 0.004 s/bp/Ma for the pre-KPg calibration age. The initial rates with the older (80 Mya) calibration (Fig. 3) are lower than those of the younger (63 Mya) calibration (Fig. 2), however the general trend is similar, with the rates near the origin of the clade being approximately ten times higher than those at the recent with a group of seeming “outlier” rates that remain high into the recent. The slope of the rates is shallower with the older calibration than with the younger calibration, but the results of both calibrations show an increase in lineages through time with a decrease in overall rates through time (Fig. 2; Fig. 3). The drastic increase in lineages matches with a higher rate of both lineage creation as well as higher nucleotide substitution rates that follow a similar trend to the LTT plot in both a pre-and post-K-Pg boundary origin scenario for the placental crown group.

When substitution rates are plotted along the tree, the highest rates along branches show a through line of higher rates from the beginning branch through to certain taxa with the highest rates being present in the rodents and other small-bodied taxa. Larger-bodied taxa have relatively low rates in comparison, and this distinction is made early in the phylogeny with a rapid split between higher and lower rates along the tree. The majority of fast rates plot along the lineages leading to the rodents as well as the soricids while the slowest rates rapidly occur along the lineages leading to primates, Boreoeutheria, and the monotreme and marsupial outgroups (Fig. 4)

**Figure 4.**
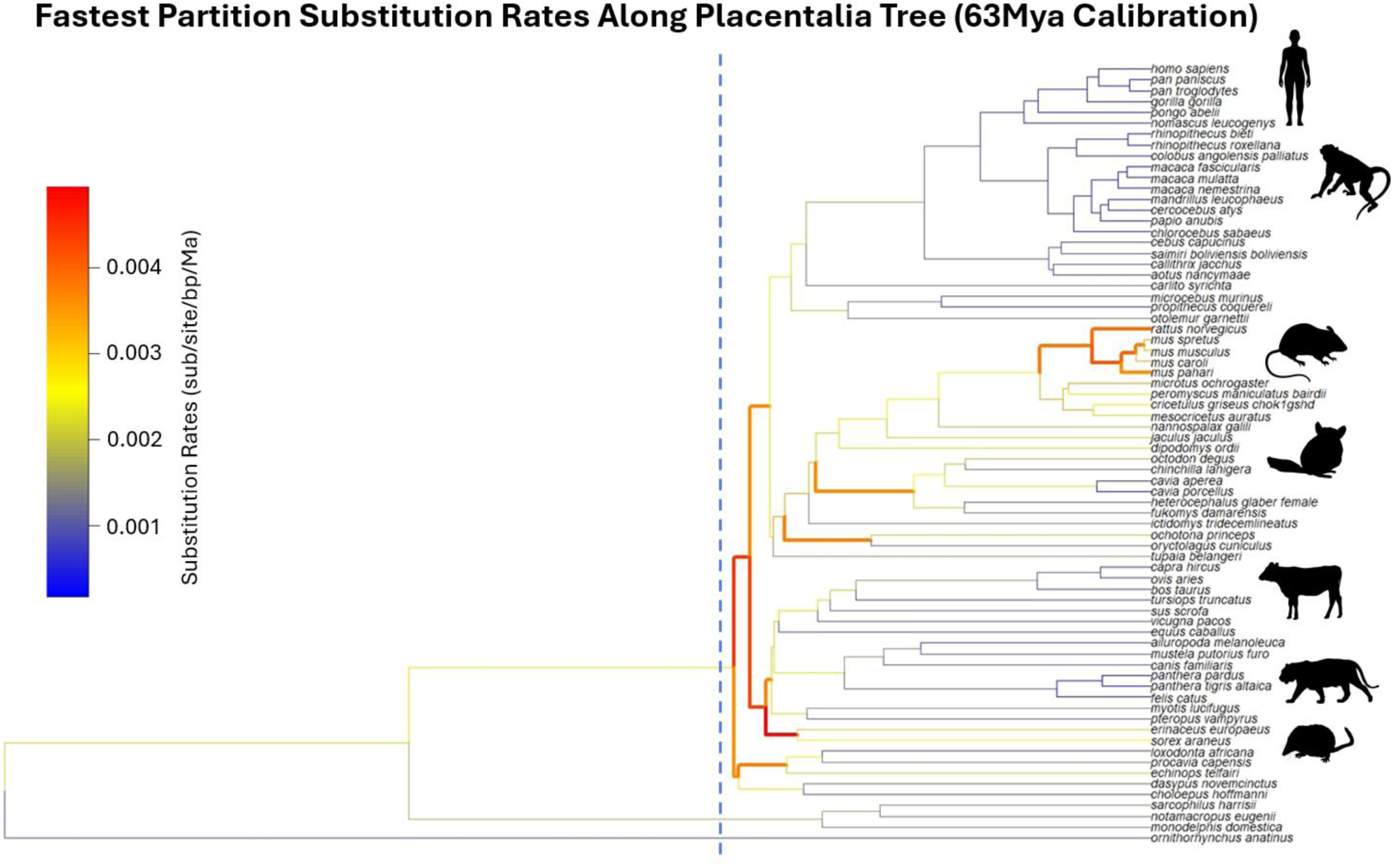
Substitution rates plotted along the Placentalia tree with highest rates in warm colours and slower rates in cool colours representing substitutions per base pair per million years. The dashed blue line indicates the K-Pg boundary at approximately 66Mya. Note that high Recent rates descend from the high early rates through lineages that also have an elevated rate. (Silhouettes adapted from Phylo Pic).

## Discussion

### Priors and posteriors

In Bayesian analysis, the posterior is proportional to the likelihood of an event multiplied by the prior. As a direct corollary, it follows that the prior always exerts at least *some* influence on the posterior, and this is true in Bayesian molecular clock analyses too (Barba-Montoya et al., 2017; Winter & Depaoli, 2023). In addition, as has been repeatedly shown, in at least some forms in which calibration information from fossil ages is inserted into clock analyses, the *effective* (i.e. joint) prior often greatly differs from the actual calibration used. Although the two are sometimes used rather interchangeably, it is important to stress that while the calibration represents the mechanics by which fossil dating information is entered into the analysis, it is only the effective prior that is worked on by the clock mechanism, and this may conflict strongly with the actual evidence from the fossil record and can lead to inaccurate estimates of clade origins, with clocks often calculating clades to be much older than the fossil record suggests.

In Bayesian analysis, it is commonly assumed or hoped that the likelihood will overwhelm the prior, so that the posterior will generally reflect the true value of the variable being estimated, no matter what the prior is. Indeed, this thought seems to lie behind the general view that the origins of placental mammals *must* lie in the Cretaceous because that is when several different clock analyses with differing methodologies have inferred it to be. For example, Benton et al. (2009) make an exception for placentals to their normal rules about fixing the maximum age of calibration ranges on precisely these grounds. In the present case however, we show that the prior continues to exert an important influence over the posterior, even when comparatively young ages are chosen for it (Fig. 1). Thus, the hope that the true age will emerge as a posterior no matter what prior is chosen is shown to be unrealistic.

We wish to stress that when the crown-group placental mammals emerged is an empirical question. Even if one accepts that crown groups are likely to emerge directly after mass extinctions, this does not rule out them emerging before. Similarly, even if no Cretaceous crown-group mammals have been found (or at least recognized) does not rule out any future such finding, or the confirmation that suggestions of crown-group membership of Cretaceous taxa (e.g. Brady et al. 2024) are correct. Indeed, a single verified Cretaceous crown-group placental mammal would be enough to falsify the hard explosive model.

### Rates of evolution

When the crown placentals are constrained to lie after the K-Pg boundary, the molecular rates of change within them reach a maximum of just above 0.006 substitutions per base pair per million years in the fastest partition and drop rapidly to about 1/10^th^ that amount by the Recent (Fig. 2). This overall trend of rapid rates decreasing approximately tenfold towards the Recent is consistent across partitions (supplemental information Fig. S3-S7). However, there are also some high rates in the Recent, with averages close to 0.004 s/bp/my for the fastest partition, and when the rates are plotted along the tree, these higher rates in the Recent fall along the branches leading to rodents and other small-bodied taxa (Fig. 4). In addition, the early stage of the inferred radiation, as represented by the lineage-through-time (LTT) plot, is also characterized by rapid diversification. In the case of the calibration at 80 Ma, rates are also at least somewhat elevated at the beginning of the crown placental radiation, even though the clock infers a crown placental origin of c. 76.6 Ma (Fig. 3). While the rates are not as high as when the calibration is set after the Cretaceous (Fig. 2), there is still a similar decreasing trend in rates towards the Recent, with one set of rates in the Recent having a roughly tenfold difference from the rates at the beginning of the clade. This consistent pattern shared between the two calibration schemes suggests that the trend in rates is not simply an artefact of forcing the clock to conform to a placental crown origin after the K-Pg boundary, but rather a general feature of the placental radiation (c.f. Phillips 2016, Phillips & Fruciano, 2018). One feature common to both our two focal analyses and that of Álvarez-Carretero (2021) is that many extant orders are given a soft maximum calibration at the K-Pg boundary, which is also bound to generate short, rapid early branches.

The possibility of an entirely post Cretaceous origin of crown-group placentals has been deprecated on the grounds of it implying exceptionally rapid, indeed “virus-like” speeds of molecular evolution at their origin (Springer et al., 2013; c.f. Álvarez-Carretero et al., 2026). When Phillips (2016) modelled the “soft explosive” origin of placentals, he recovered somewhat faster rates at the beginning of the radiation (his fig. 5), with rates ranging from about 0.004 s/bp/Myrs to about 0.0007 s/bp/Myrs near the present (note that the scale in the figure should be per hundred million, not million, years), but nothing as fast as that implied by Springer et al. (2013). In our analysis we recovered faster rates than those of Phillips in the fast partition, as expected; in the combined partitions our rates were similar to his.

The higher rates at the beginning of the clade according to our analysis, at c. 0.006 s/bp/Myrs, although high, fall within the range of substitution rates present in modern rodent taxa, and averaging across partitions (supplemental information Fig. S7), these same rates fall within the average rates for mammalian genes across the entire clade (Bininda-Emonds, 2007; Kumar & Subramanian, 2002) Thus, like Phillips (2016) and O’Leary et al. (2013b), we recover rates at the origin of the clade to be within the gamut of recent placentals (c.f. Bergeron et al., 2023 who recover a comparable range of generational mutation rates). The other striking feature of the post-boundary calibration is that the high rates at the origin are linked via intervening lineages to the high rates today. This implies that high rates are inherited along the lineages.

In vertebrates in general, high mutation rates have been linked to small body size, high generation rates, and high metabolic rates (Li et al., 1996; Martin & Palumbi, 1993; Wu & Li, 1985; but see also Liow et al., 2017). This would particularly be the case if the last common crown-placental ancestor was a small, rodent-like animal as suggested by hypotheses for the crown-placental ancestor such as that of O’Leary et al. (2013a) which posited that the common placental ancestor was a rodent-like insectivore with a small body size, fast metabolism, and thus likely a relatively fast generation time similar to modern species such as mice or shrews.

### Placental evolution and the Covariant Evolutionary Tempo model

Despite the known relationship between body size and evolutionary rates in mammals, few studies have attempted to connect these variables to molecular clock analyses, with notable exceptions being Springer et al. (2003), Bromham (2011); Phillips (2016), and Phillips and Fruciano (2018). However, as noted especially by Bromham (2011), systematic body size differences across the clade are likely to significantly impact the results of molecular clock studies. Similar remarks were made by Beck and Lee (2014) when examining a morphological clock model, and both studies suggest that the data are compatible with a post K-Pg origin of placentals. These observations fit the newly-developed “Covariant Evolutionary Tempo” (CET) model (Budd & Mann, 2025), which posits that molecular (and potentially morphological) evolutionary and diversification rates covary through time. The outcome of this model is that evolutionary rates in major clades would tend to be characterised by short early periods of rapid diversification and molecular change, which in general should decay to slower rates, although elevated rates should persist along some lineages, so that the spread of recent rates should contain examples comparable to the high early rates. Another prediction is that clade sizes of sister groups should be highly asymmetrical, with diversity being dominated by one of the two. Indeed, it is notable that in a typical phylogeny of extant placentals, the Atlantogenata-Boreoeutheria split has a species split that is approximately 120 to 5000+, the sort of clade imbalance typical of CET outcomes, and extremely implausible under homogeneous birth death processes or purely secular variation.

We have shown in this paper that the posterior of crown-group placental origins depends on the maximum age of the calibration, and that post-Cretaceous origins can be recovered with soft maximum calibration ages a little younger than the boundary. This might suggest that the molecular clock itself cannot distinguish between a pre-and post-boundary origin, as the result depends heavily on the calibration and thus prior choice. However, a recent paper (Álvarez-Carretero et al. 2026) attempts to resolve this problem via Bayesian model selection, in order to distinguish which of the two origins is most likely, and conclude that the post-Cretaceous origin is vanishingly unlikely. However, the validity of this analysis depends on the underlying model from which likelihoods are extracted being true. In particular, the combination of very short branches and very fast rates would be extremely unlikely under their model. This is because of two factors. The first of these is that their model does not allow for any variation in diversification parameters (e.g. the ClaDS model of Maliet et al., 2019, Barido-Sottani & Morlon, 2023, and the birth-death diffusion model of Quintero et al., 2024), which inherently disfavours rapid radiations. The second is that diversification and molecular evolution are modelled as independent processes; this implies that it would be very unlikely for both to be elevated simultaneously. If this model is wrong, as we and others have argued here and elsewhere (Hua & Bromham, 2017; Ritchie et al., 2022; Budd & Mann, 2025), then Bayesian model selection within its scope will simply mislead. The hypothesis of an extremely rapid radiation of placental mammals (or other clades) is in this light seen to be strongly discriminated against in most clock models. Such radiations have been recovered by other recent authors, such as Carlisle et al., 2023 for metazoans and Wu et al., 2026 for angiosperms; presumably, these inferred rapid radiations would also be found to have negligible posterior support under a Bayesian model selection.

### The plausibility of a rapid radiation

The description of *Purgatorius* spp. from less than 140 Kyrs above the K-Pg boundary in Montana (Wilson Mantilla et al., 2021) might suggest that definitive crown-group placentals (i.e. stem-group primates) are present almost immediately after the mass extinction, which to many will suggest that an origin before the boundary is almost inevitable. However, there are at least two considerations that might count against this. The first is that the phylogenetic placement of many basal forms in a phylogeny is problematic, and that is true of *Purgatorius* too. After all, the claim that early crown-group forms closely resembled the late stem-group forms, so that we might be misidentifying Cretaceous crown-group taxa as stem group ones surely must cut both ways, and allow the possibility that apparent basal crown-group forms in the Paleogene are misidentified stem-group members. Secondly, if these very early forms do turn out to be crown-group placentals, these particular fossils would imply perhaps five or so lineages emerging within less than 150 Kyrs. This might seem straightforwardly impossible. However, there is no reason to think that a speciation that in the end leads to a major clade such as an order should be any different from any other speciation event, so there is no reason for it to in some way “take more time”. At the beginning of what in retrospect is going to be a major clade, most if not all speciation events will lead to lineages and not plesions (c.f. Budd & Mann, 2018; c.f. Helmstetter et al., 2022; Pennell et al., 2012). Indeed, a radiation of this sort might be comparable to that of, for example, the cichlids in Lake Victoria (Nakamura et al., 2021) where perhaps 500 species are known to have arisen within <15 Kyrs. Set in this context, the origin of the placental mammals might have been exceptionally rapid, but still well within the limits of known rapid radiations.

Finally, we note that the patterns of rapid early diversification and molecular change we have observed here are not confined to the placental mammals – similar patterns have been recorded in birds (Berv et al., 2018) and arthropods (Lee et al., 2013), supporting the idea that these patterns may be a more general feature of major radiations.

## Supplemental Information

**Table S1.**
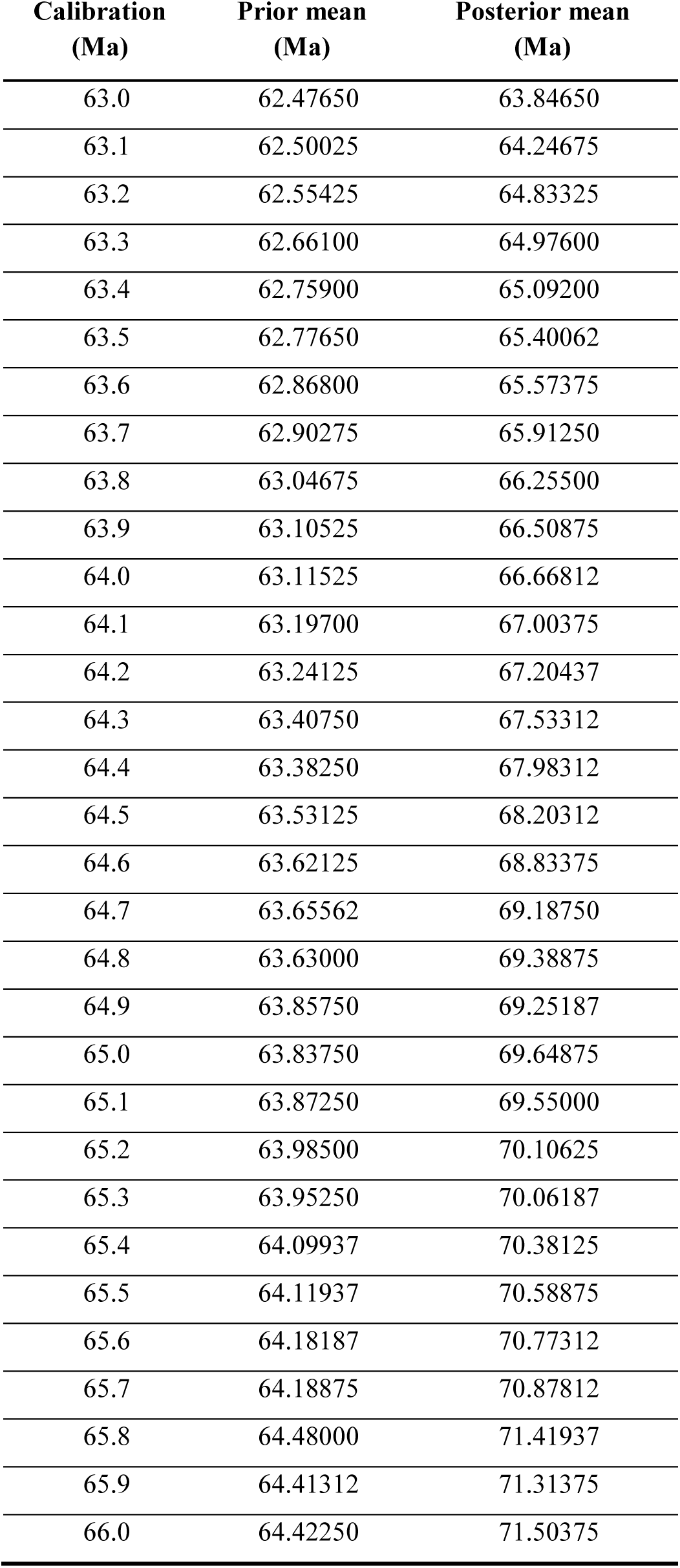
Prior and posterior mean estimates for the base of Placentalia at each calibration age every 100ka from 63Ma to 66Ma.

| **Calibration**<br>**(Ma)** | **Prior mean**<br>**(Ma)** | **Posterior mean**<br>**(Ma)** |
| --- | --- | --- |
| 63.0 | 62.47650 | 63.84650 |
| 63.1 | 62.50025 | 64.24675 |
| 63.2 | 62.55425 | 64.83325 |
| 63.3 | 62.66100 | 64.97600 |
| 63.4 | 62.75900 | 65.09200 |
| 63.5 | 62.77650 | 65.40062 |
| 63.6 | 62.86800 | 65.57375 |
| 63.7 | 62.90275 | 65.91250 |
| 63.8 | 63.04675 | 66.25500 |
| 63.9 | 63.10525 | 66.50875 |
| 64.0 | 63.11525 | 66.66812 |
| 64.1 | 63.19700 | 67.00375 |
| 64.2 | 63.24125 | 67.20437 |
| 64.3 | 63.40750 | 67.53312 |
| 64.4 | 63.38250 | 67.98312 |
| 64.5 | 63.53125 | 68.20312 |
| 64.6 | 63.62125 | 68.83375 |
| 64.7 | 63.65562 | 69.18750 |
| 64.8 | 63.63000 | 69.38875 |
| 64.9 | 63.85750 | 69.25187 |
| 65.0 | 63.83750 | 69.64875 |
| 65.1 | 63.87250 | 69.55000 |
| 65.2 | 63.98500 | 70.10625 |
| 65.3 | 63.95250 | 70.06187 |
| 65.4 | 64.09937 | 70.38125 |
| 65.5 | 64.11937 | 70.58875 |
| 65.6 | 64.18187 | 70.77312 |
| 65.7 | 64.18875 | 70.87812 |
| 65.8 | 64.48000 | 71.41937 |
| 65.9 | 64.41312 | 71.31375 |
| 66.0 | 64.42250 | 71.50375 |

---

**Figure S1.**
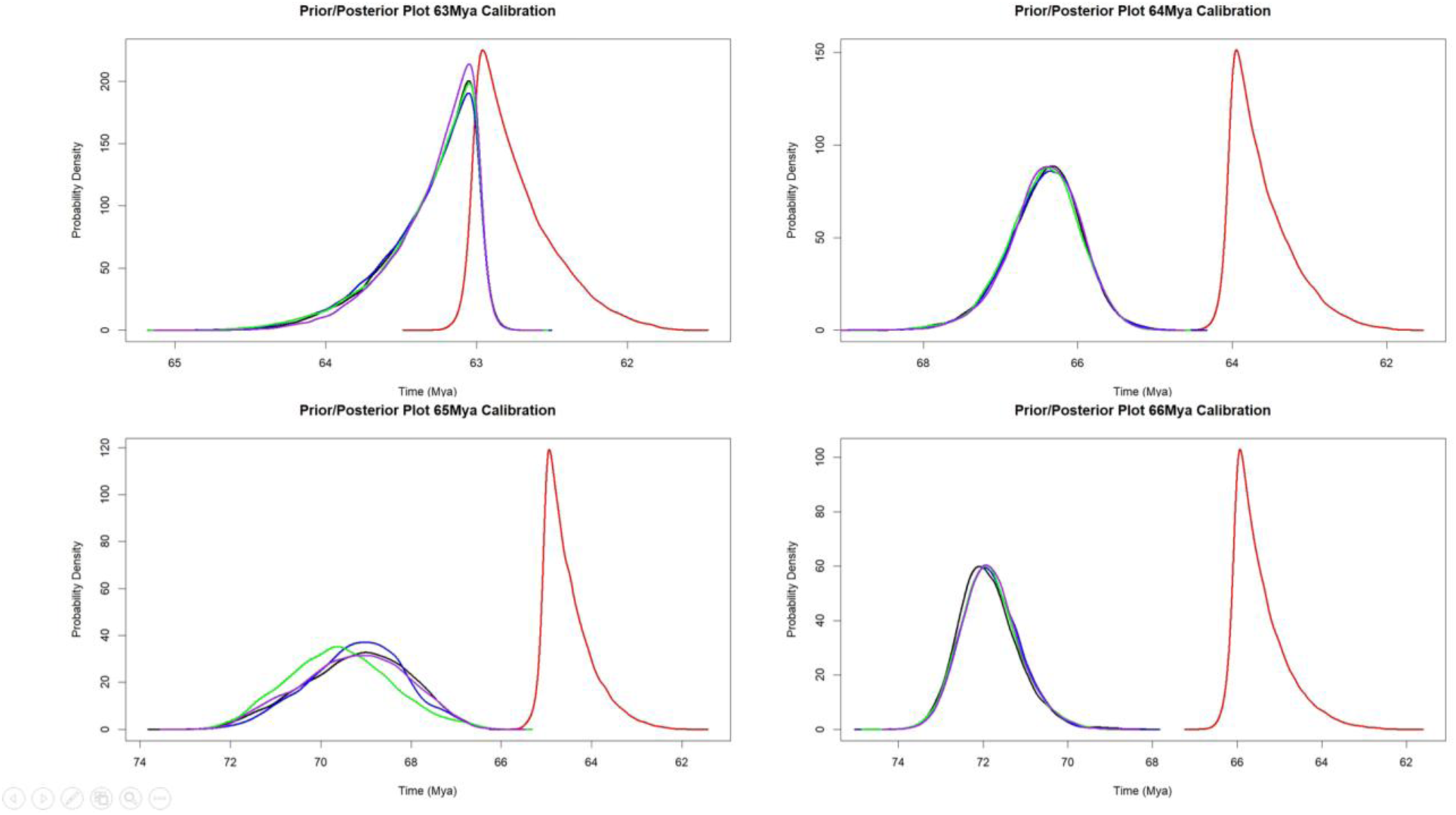
Prior-Posterior plots of four test calibrations from 63-66Mya for the base of Placentalia. Priors are in red and four separate runs for the posterior are represented in black, purple, green, and blue respectively. Increasing the age of the base calibration results in a decrease in the overlap between the prior and posterior.

**Figure S2.**
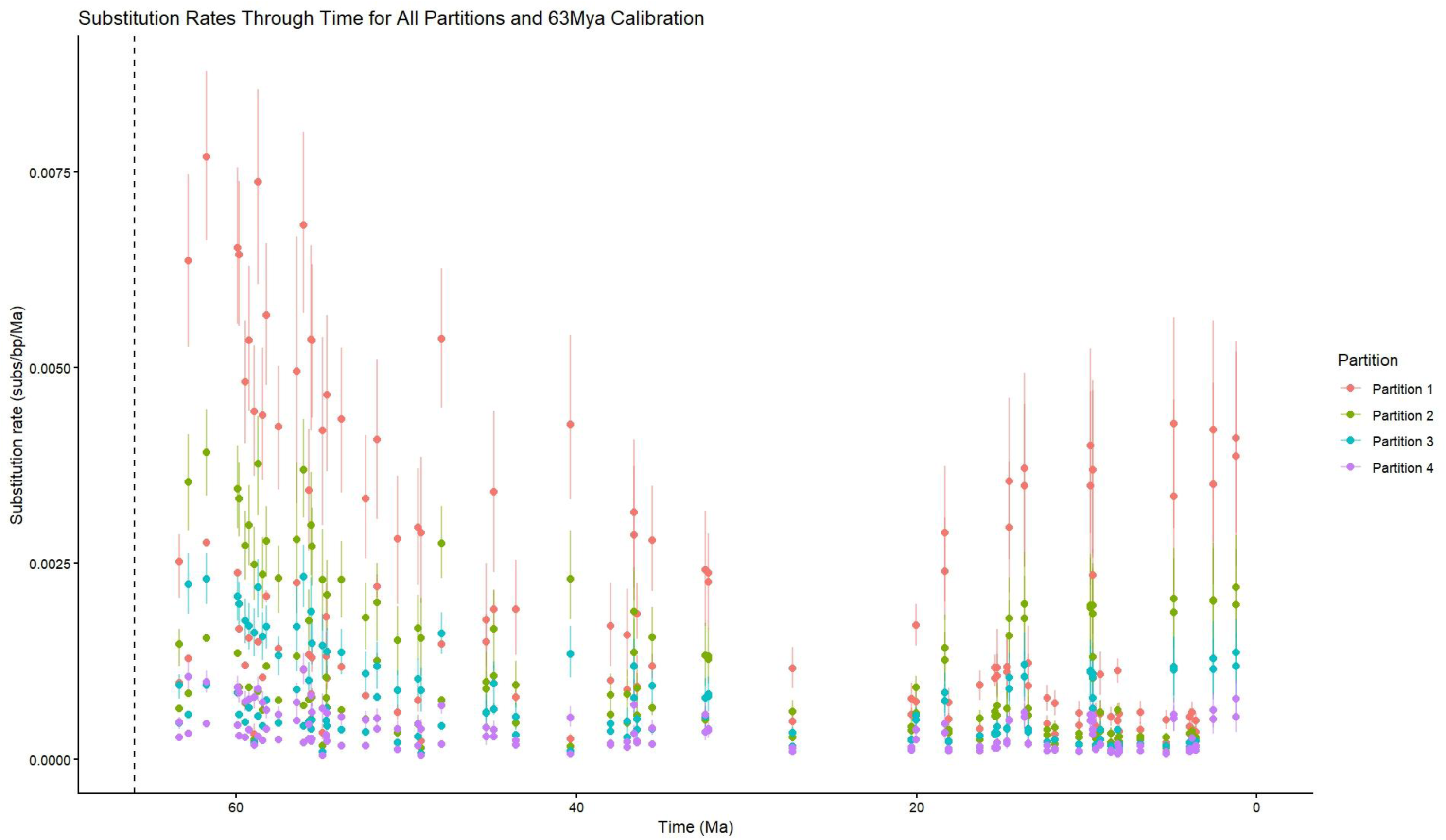
Plot of rates and 95% confidence interval for each of four partitions across time. Partition 1 is the fastest partition and partition 4 is the slowest. The vertical dashed line denotes the KPg boundary at 66Mya.

**Figure S3.**
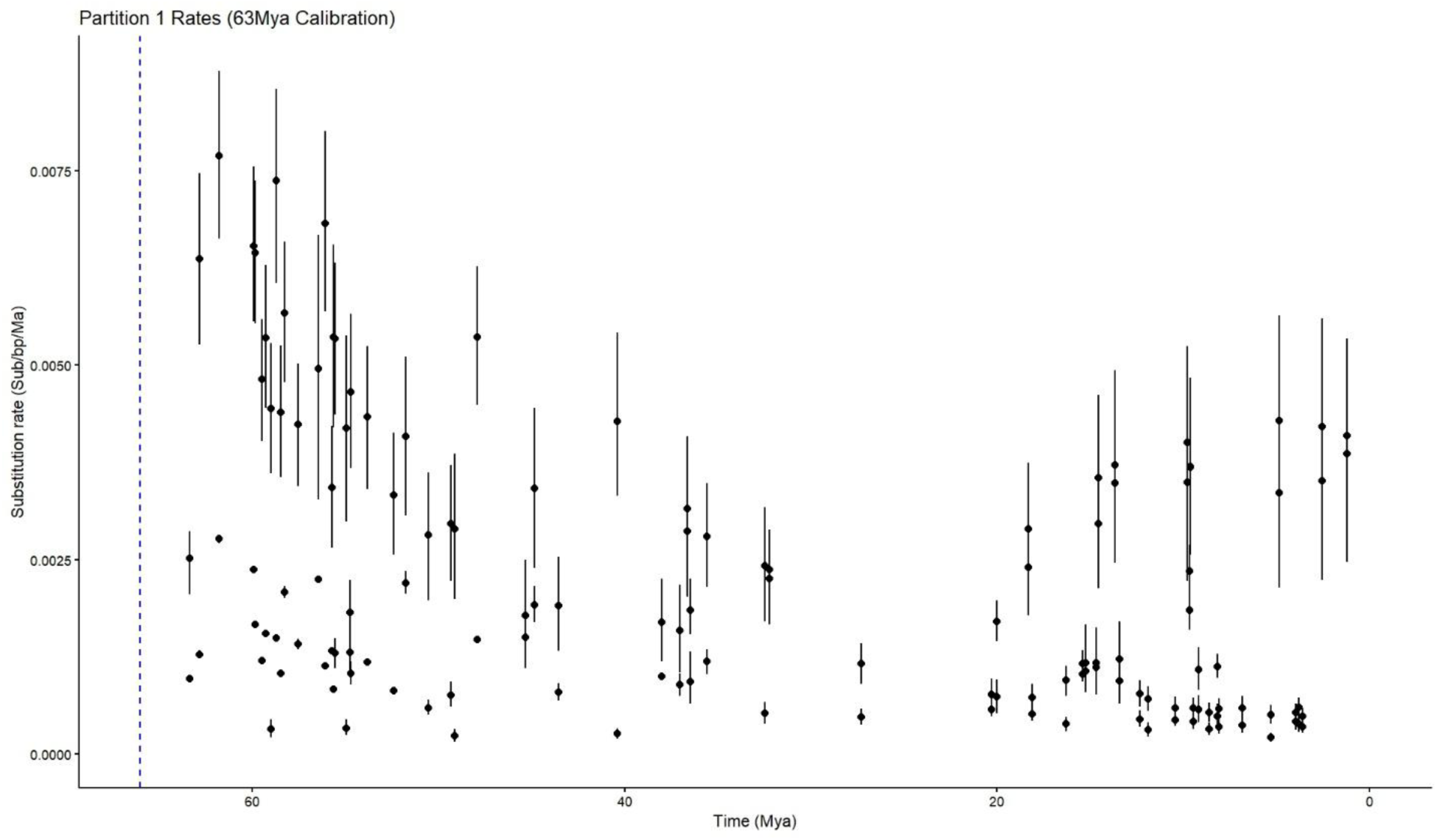
Rates through time for the fastest partition (partition 1) with 95% confidence intervals. Blue dashed line denotes the K-Pg boundary at 66Mya.

**Figure S4.**
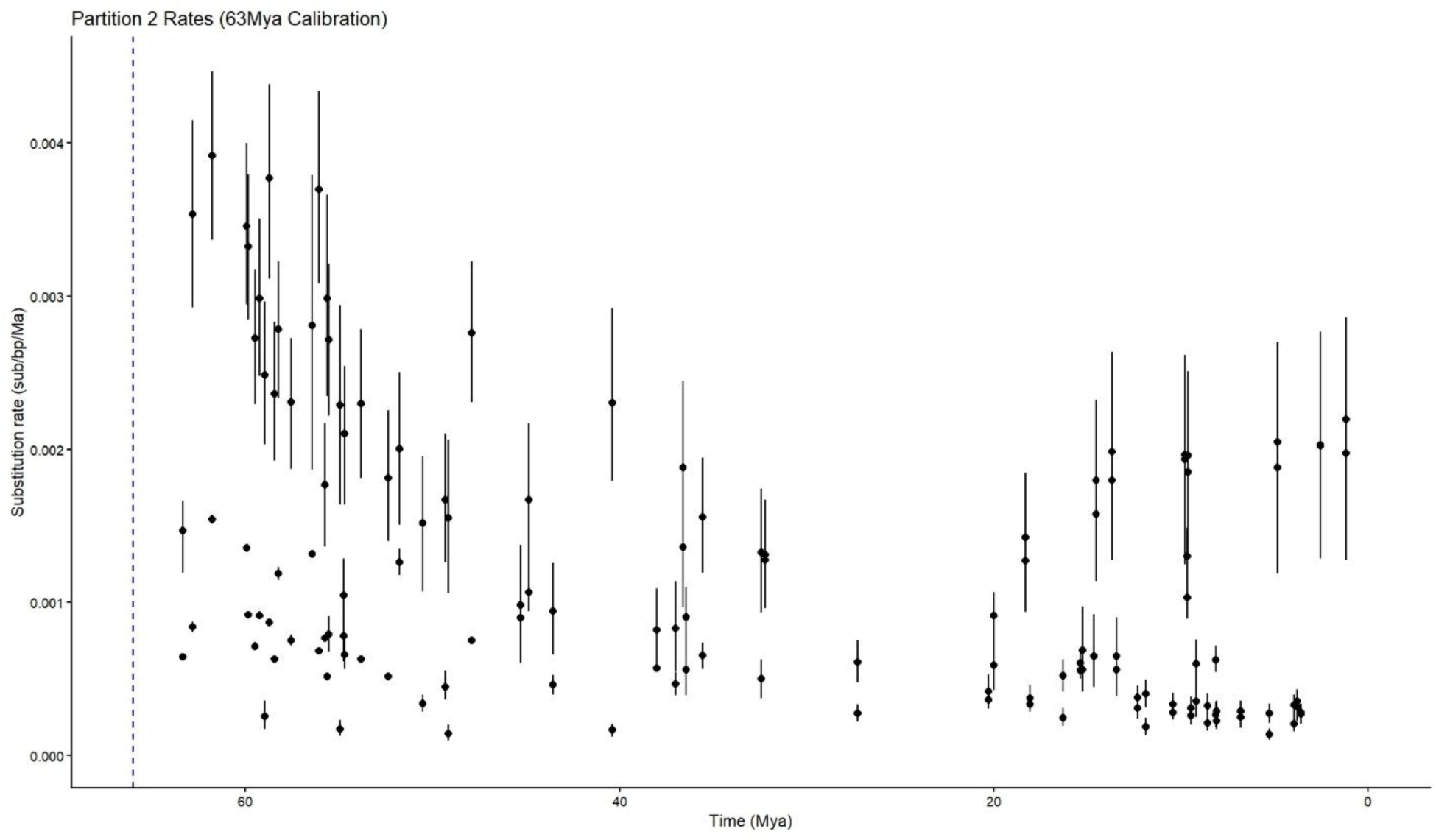
Rates through time for the second fastest partition (partition 2) with 95% confidence intervals. Blue dashed line denotes the K-Pg boundary at 66Mya.

**Figure S5.**
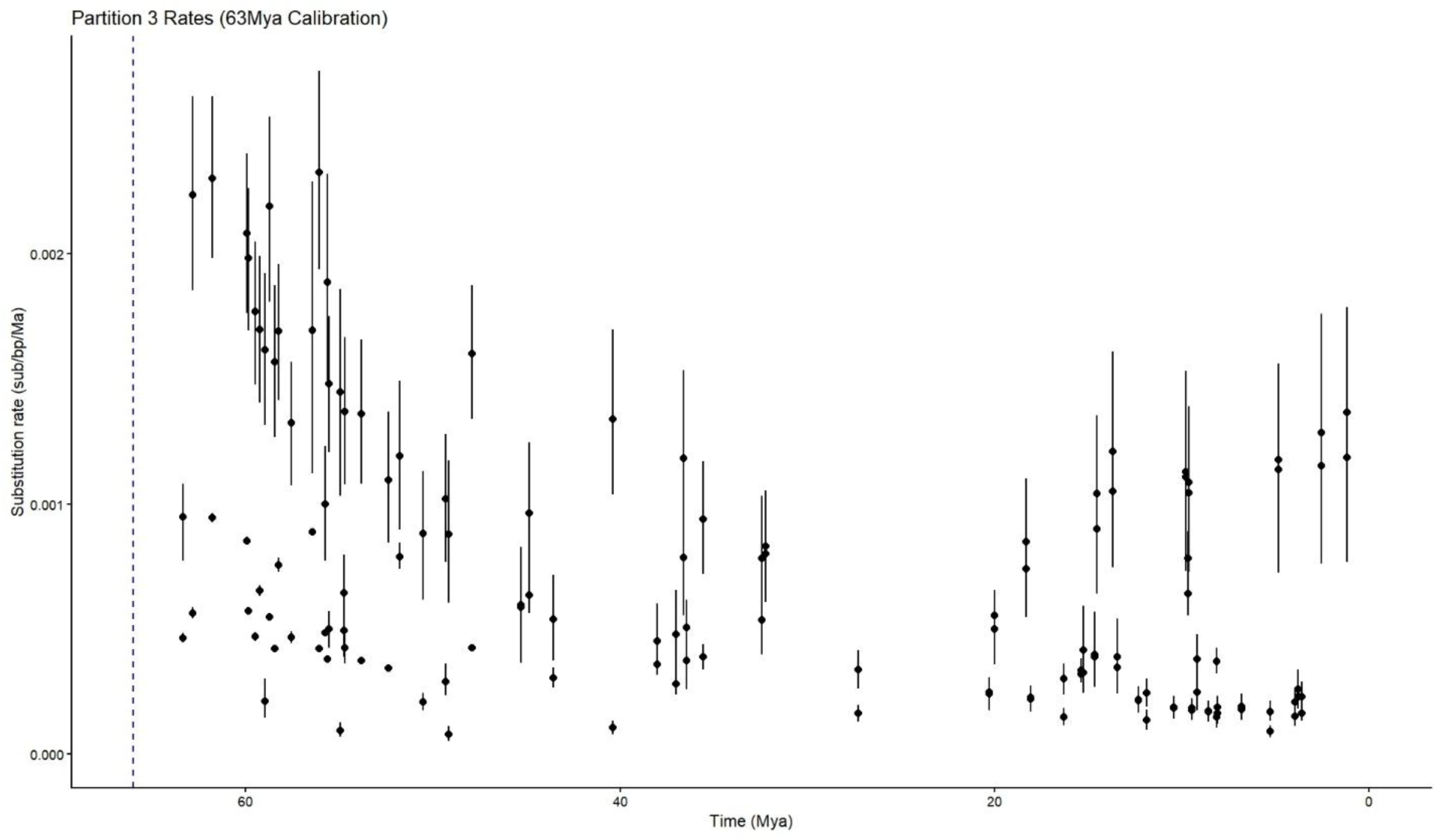
Rates through time for partition 3 with 95% confidence intervals. Blue dashed line denotes the K-Pg boundary at 66Mya.

**Figure S6.**
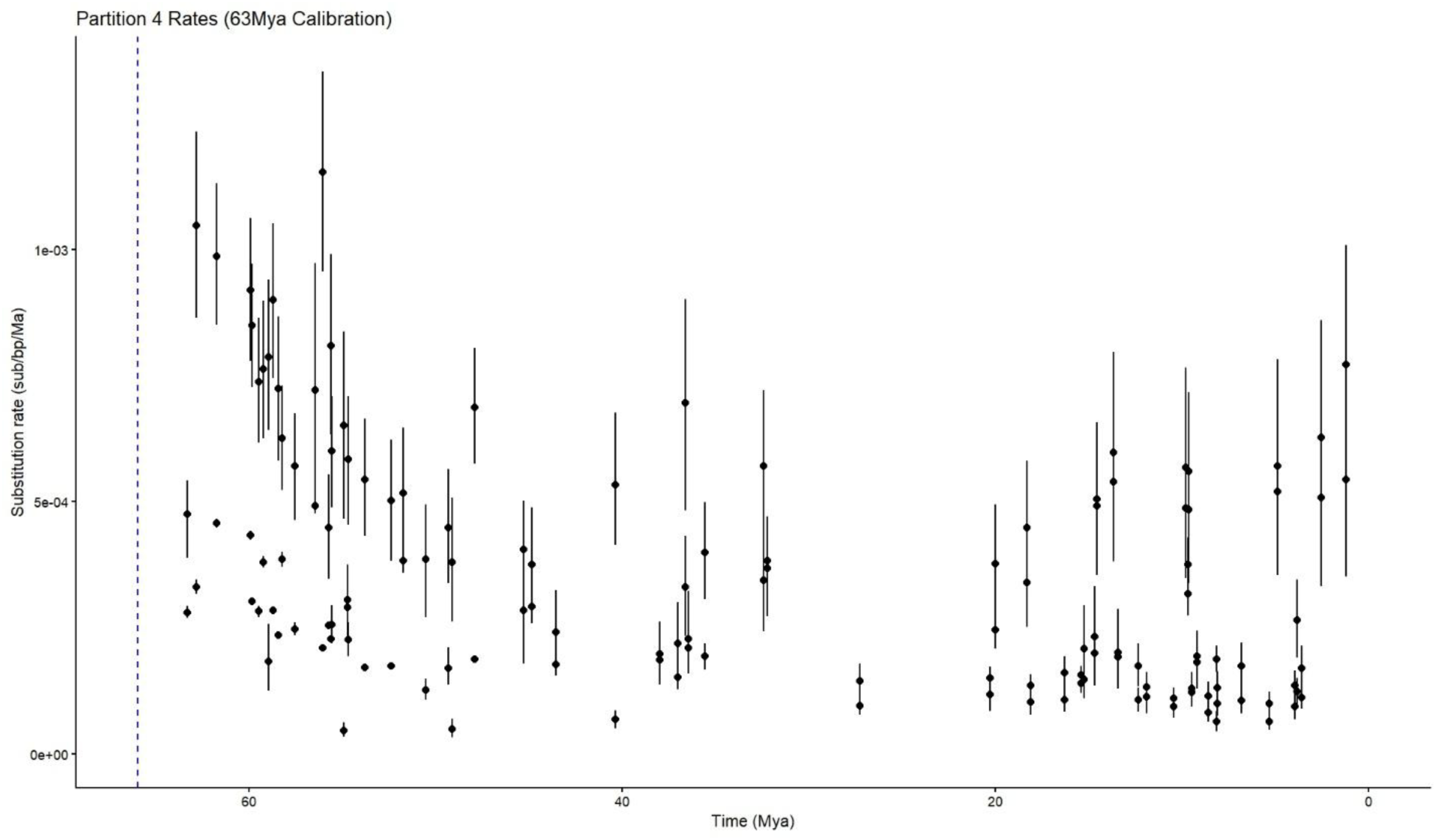
Rates through time for the slowest partition (partition 4) with 95% confidence intervals. Blue dashed line denotes the K-Pg boundary at 66Mya.

**Figure S7.**
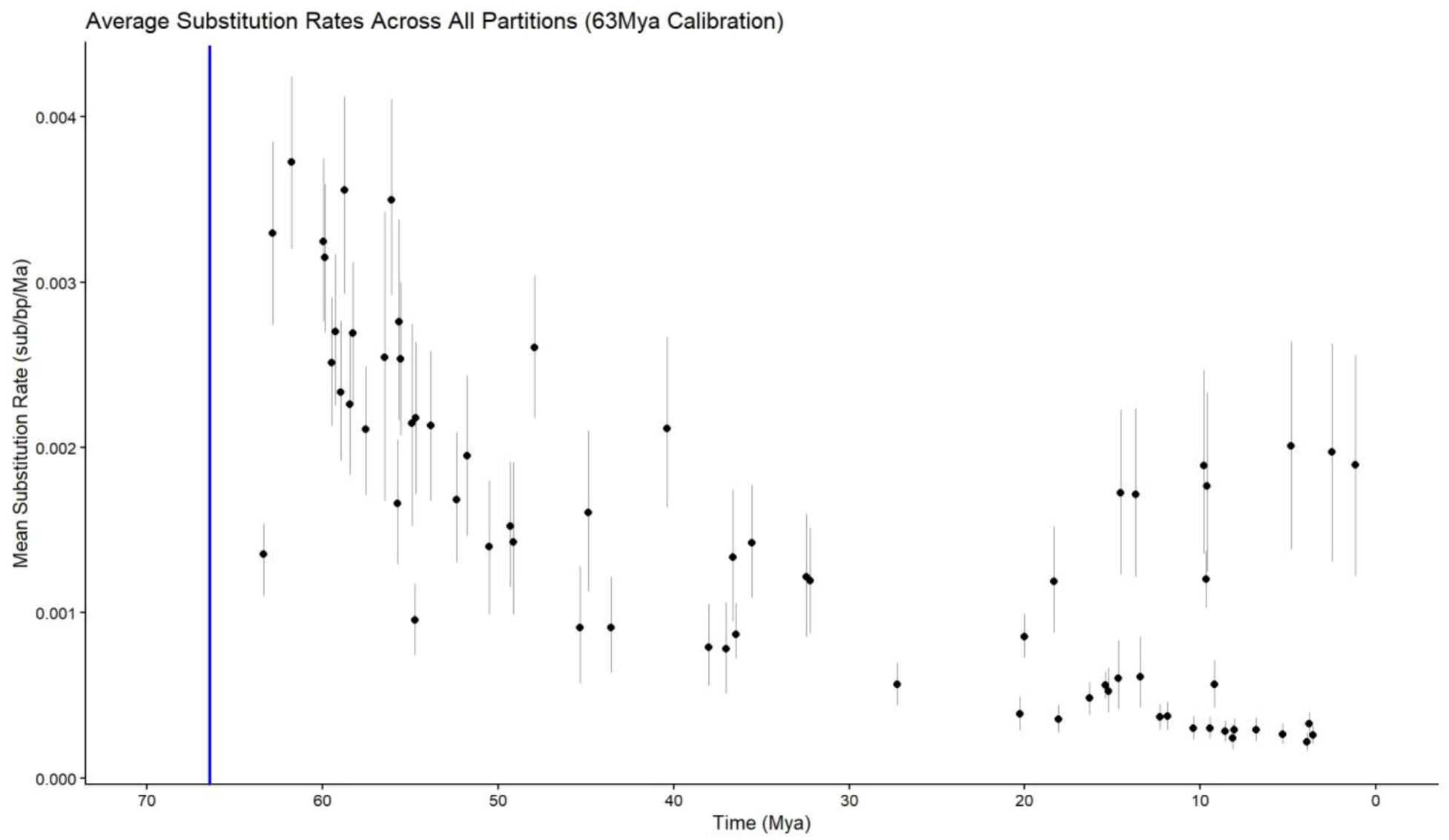
Rates through time averaged across all partitions with 95% confidence intervals. Blue line denotes the K-Pg boundary at 66Mya.

